# Designing and validating a staircase to leverage floor mounted force plates

**DOI:** 10.1101/718346

**Authors:** Todd J. Hullfish, Josh R. Baxter

## Abstract

Navigating stairs is a challenging task for many patient populations. Unfortunately, assessing lower extremity kinetics is not practical in many laboratories due in part to methodologic constraints. In this study, we designed, fabricated, and calibrated a staircase that accurately measured ground reaction forces applied to the second and fourth step. This implementation met several design criteria that included low-cost, ability to quickly move the staircase in and out of motion capture spaces, stable and safe staircase, and easily modifiable to meet the constraints of different lab layouts. We built the staircase as an outer and inner staircase assembly constructed using a modular aluminum framing system. Once positioned on our force plates that were embedded in the lab floor, we used an instrumented pole to apply known loads to a series of surface locations on the force plates and steps that were resting on top of the force plates. This calibration procedure reduced the center of pressure errors by approximately 50% for the embedded force plates and lower step (step 2) and 3-fold for the higher step (step 4). Next, we demonstrated that these steps can be integrated into a clinical gait analysis workflow. A single healthy-young adult navigated the stairs, the ground reaction forces were transformed into stair reaction forces, and these external loads were used to solve the inverse dynamics problem. This staircase provides other researchers with a new tool to assess stair navigation biomechanics. In this study, we provided the bill of materials, mechanical drawings, and calibration code necessary to modify and implement this staircase paradigm into other lab layouts.

## Introduction

Navigating stairs is an activity of daily living that presents functional challenges and injury risks for elderly adults [1] and many patients with musculoskeletal [2] and neurologic [3] pathologies. Nearly one-million fall injuries occur annually during stair navigation, accounting for 14% of all fall injuries [4]. Unlike walking over flat ground that is a simple activity for many populations, ascending and descending stairs requires increased muscle shortening [5] and joint motion and kinetics [6] to safely complete these tasks. Navigating stairs becomes more challenging with increased body mass and stair height [7], highlighting the importance of understanding the biomechanics of stair navigation when planning for safe practices in special populations.

While analyzing stair navigation biomechanics provide important insight into ambulatory function and musculoskeletal loading [8], logistical constraints prevent stair navigation from being integrated into many traditional gait analysis laboratories. Instrumented staircases have traditionally required rigid fixation to ground mounted force plates [9], which cannot easily be achieved after initial force plate installation, or expensive force plates embedded in each step [5,10]. These instrumented staircases are also heavy and not easily moved, placing additional constraints on experimental design by limiting the number of activities that can be analyzed during a single patient visit. As many laboratories are designed to acquire motion capture data centered around pre-installed force plates, staircases that are either permanently installed or require lengthy installations onto force plates increases the time required to perform analyses.

The purpose of this study was to develop and validate a custom built staircase. During the development of this staircase, we determined that several design criteria were necessary to ensure the feasibility of implementing the staircase in biomechanical research: 1) the staircase must be low-cost, we defined this as under $5,000 US; 2) the staircase must be mobile and operate in motion capture volumes without the need to move or recalibrate cameras; 3) the staircase must be safe to use, both in terms of its general stability and general dimensions; and 4) the staircase must be easy to manufacture and implement in other research labs. Once the staircase was developed, we implemented a calibration paradigm that has been reported previously to calibrate both force plates and instrumented treadmills [11]. Finally, we determined the effects of force plate and stair calibration on ankle, knee, and hip joint moments calculated using inverse dynamics.

## Methods

### Study Overview

To develop and validate a custom built staircase, we performed three discrete tasks. First, we designed and fabricated a staircase based on our design criteria. Based on the physical constraints of the force plates in our laboratory, we designed a staircase with 4 steps and a landing. The second and fourth steps were mechanically isolated from the staircase assembly and rested on the three laboratory force plates. Second, we used an instrumented pole to calibrate and validate the force-moment measurements of each force plate and the two steps that were resting on the force plates. Third, we analyzed gait biomechanics during walking over flat ground and stair navigation to determine the fidelity of joint kinetics calculated with this approach. As part of this study, we have provided a complete bill of materials for the described staircase and analysis code for the calibration and validation of the staircase.

### Staircase Design and Fabrication

To satisfy our design criteria that the staircase be mobile, low-cost, stable and safe, and easy to manufacture; we decided to use aluminum t-slot structural framing that was cut to length with fastener cutouts performed directly from the vendor (80/20 Inc., Columbia City, Indiana, USA). The staircase (**Figure 1**) was comprised of 4 steps (0.76 m wide, 0.31 m deep,0.197 m high) and a top landing area (1.06 m wide, 0.89 m deep) that conformed with federal guidelines for stairs (Occupational Safety and Health Administration, USA). We installed handrails on both sides of the staircase and optional support struts to increase the rotational stability of the staircase assembly in the event of patients losing their balance or falling. The structural framing used in this staircase assembly cost less than our threshold for ‘low-cost’ (Total US $3,450 in 2018, bill of materials available in supplemental material).

**Figure 1.**
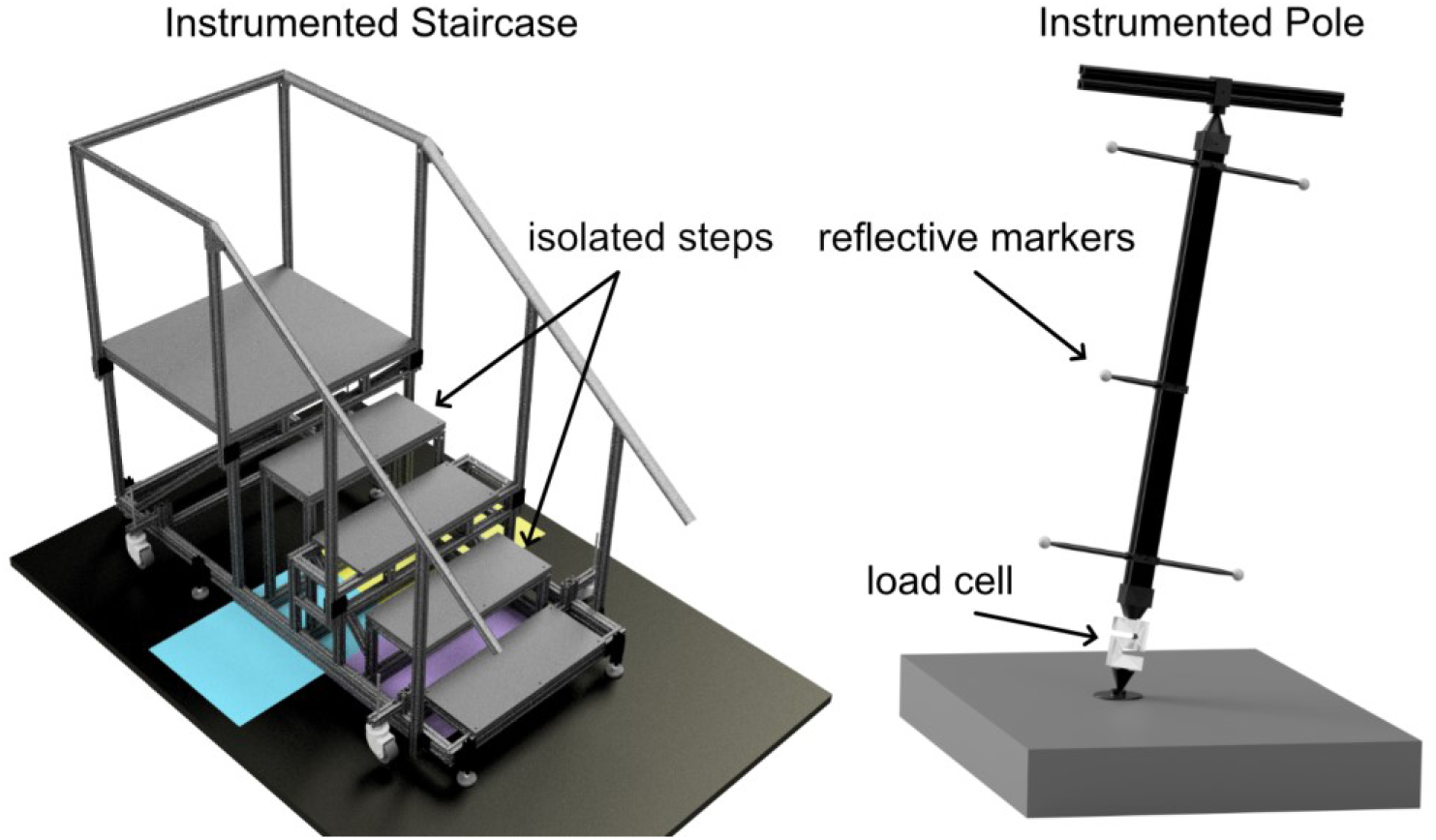
(1.5 columns). We fabricated an instrumented staircase (*left*) using a modular aluminum framing system. This staircase had two main assemblies: an outer assembly that rested on the laboratory floor and consisted of the first and third steps as well as the top landing and hand rails and an inner assembly that rested on the three force plates (*color rectangles*) that were embedded in the lab floor and consisted of the second and fourth steps. We calibrated these force plates and steps by applying known loads using an instrumented pole (*right*) and a least squares approach described previously in the literature [11].

The staircase was built as two separate structures: the outer staircase assembly, which was comprised of the first and third steps and the top landing; and the inner staircase assembly, which was comprised of the second (0.395 m high) and fourth (0.795 m high) steps. The outer staircase assembly rested on the laboratory floor using six self-leveling feet. The inner staircase assembly was mechanically isolated from the outer staircase assembly and rested fully on the three laboratory ground mounted force plates (BP600900, AMTI, Watertown Massachusetts, USA) that were arranged in a pyramid layout (**Figure 1**). To increase the rigidity of the inner staircase assembly, we used 45-degree support assemblies to connect the low step (second step) with the high step (fourth step). During development, we used hard rubber feet on the four corners of the inner staircase assembly but found that this caused an appreciable amount of movement of the step during testing. Our force plates have a thin rubber matting (3mm thick), which provided a small amount of compliance that allowed for continuous contact between the length of the structural framing (1.20 m long) and the force plates.

To minimize the setup time and make flat ground and stair navigation analyses in a single study session practical in a busy clinical gait analysis setting, we designed the staircase assembly with a caster system that is engaged by driving a lead screw using a cordless power drill. However, there are other commercially available solutions for moving large structures like workbenches that would also work. To move the inner staircase assembly with the outer staircase assembly, we designed a simple support system that used two rubber coated hooks and a small piece of dimensional lumbar (2”×8” ×8”) to hold the inner staircase assembly in place while the staircase was positioned. During development and pilot testing, we were able to consistently position the staircase assembly in under two minutes.

### Calibration and Validation

To calibrate and validate our custom-built staircase, we implemented a previously reported calibration paradigm [11] to calibrate both our laboratory force plates and the staircase assembly (**Figure 1**). This paradigm used an instrumented pole (**Figure 1**, complete drawings available in supplemental material) to apply known loads on the force plates and inner staircase assembly to develop a correction matrix that would reduce the measurement errors of the force plates. Briefly, we modified a 0.72 m length of structural framing with an in-line load-cell on the distal end and hard cones on both ends of the pole, which allowed for pure axial loads to be applied to the surface being calibrated via a protective plate with a socket [11]. The load-cell (TAS501, HTC-Sensor, Xi’an, China) had a 200kg capacity and was secured to the pole with threaded steel rod. We used a microcontroller (Metro Mini, Adafruit, New York, New York, USA) and a load cell amplifier (HX711, Sparkfun, Niwot, Colorado, USA) to process the load-cell signal and output the analog signal using a 12-bit digital to analog converter (MCP4725, Adafruit, New York, New York, USA) that was synchronously logged with motion capture data on our motion capture system. A simple handle was built using a shorter 46 cm length of structural framing and a socket to accept the articulating cone that was secured to the pole. We placed five reflective markers on the pole using 3D printed marker wands and acquired the marker trajectories to calculate the direction and position of the force applied by the pole to the structure being calibrated. Motion capture data were acquired using a 12-camera motion capture system (Raptor Series, Motion Analysis Corporation, Rohnert Park, California, USA).

To calibrate the laboratory force plates, we applied known loads to each force plate and calculated a calibration matrix to correct for measurement errors. First, a single investigator loaded each surface by pressing down and rotating the instrumented pole through arcs of approximately 15 degrees (**Figure 1**). These loads were applied to twenty points evenly distributed across the length of the surfaces (**Figure 2**). Next, we imported the instrumented pole load-cell and marker data and force plate data into a scientific computing package (MATLAB 2018a, The Mathworks, Natick, MA) and transformed these signals into the lab coordinate system (See supplemental material for MATLAB processing scripts). All load and marker trajectory data were filtered using a 4^th^ order Butterworth filter with a cutoff frequency of 6 Hz [12]. Using the instrumented pole data as the ‘gold-standard’ load applied to the force plates, we used a ‘Post-Installation Least-Squares’ procedure previously described by Collins et al. [11]. Briefly, the three force and three moment vectors were concatenated into a single 6 x *n* matrix (where *n* is the number of frames where appreciable loads were applied) for the instrumented pole (denoted as *R*) and the force plate (denoted as *S*). Based on our pilot testing and other reports, we decided to calibrate the force plates and stairs with loads greater than 100 N in magnitude [11,13]. In this linearized system, the instrumented pole load data was equal to a calibration matrix multiplied by the force plate load data (*Eq* 1). This process was repeated to calculate unique *C* matrix for each force plate.

**Figure 2.**
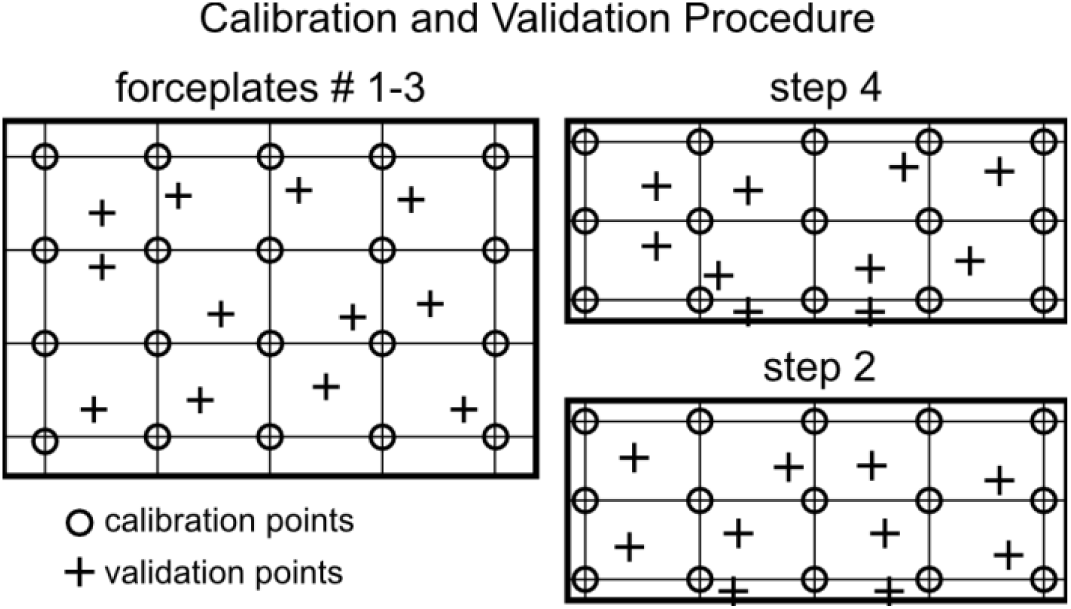
(1 column). We calibrated the 3 force plates embedded in our lab floor (*left*) and the low step (*step 2*) and high step (*step 4*) by applying known loads to a grid of 20 evenly distributed points on the force plates and 15 evenly distributed points on the steps (*circles*). Using a previously reported least squares approach [11], we calculated a calibration matrix to correct for errors in the ground reaction forces. Next, we applied known loads to randomly located validation points (*pluses*) in between each calibration point.

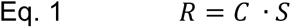

Where *C* is a 6 × 6 calibration matrix. To calculate *C*, we performed a matrix pseudo-inverse (*Eq* 2):

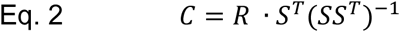

The low and high steps of the inner staircase assembly were calibrated in a similar fashion. We applied a series of fifteen loads to each of the two steps of the inner staircase assembly with the instrumented pole (**Figure 3**). Because the inner staircase assembly was positioned over the three laboratory force plates, the load data from each force plate were transformed to the lab coordinate system and combined into a single 6 x *n* matrix. We used this process to calculate a unique *C* matrix for the low and high steps.

**Figure 3.**
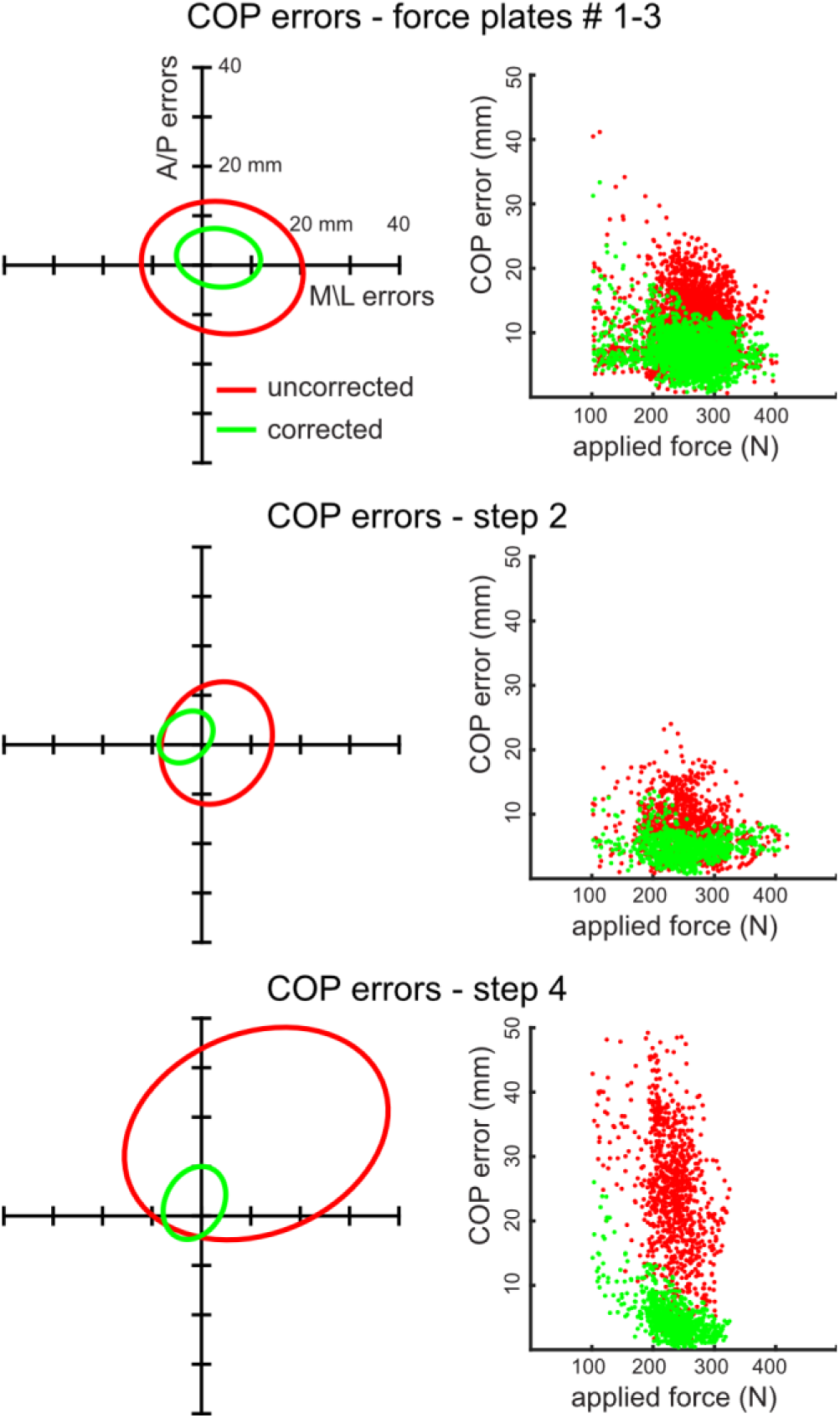
(1 column). Center of pressure errors were greater when calculated from uncorrected force plate and step measurements (*red*) than compared to corrected measurements (*green*) following the calibration procedure. The embedded force plates (*top row*) and low step (step 2 – *middle row*) produced smaller errors than the higher step (step 4 – *bottom row*). While the center of pressure errors were mostly symmetrical in both the anterior-posterior and medial-lateral directions (*left column*) for the force plates and low step, the higher step produced greater errors that were mostly anterior (the ascending direction) and medial directed. These center of pressure errors decreased with greater applied loads to the force plates and steps (*right column*).

Following the calibration procedures for the force plates and inner staircase assembly, we validated these calibrations by applying loads between each of the calibration locations on the three force plates and two steps (**Figure 2**). We calculated the root mean square error (RMSE) and 95% confidence ellipses by comparing the un-calibrated and calibrated force plate and stair centers of pressure compared to the instrumented pole measurements. To determine if load magnitude had an effect on center of pressure errors, we visualized the magnitude of loading to the errors between the instrumented pole data and the calculated data.

### Gait and Stair Navigation

To establish the efficacy of implementing this novel staircase assembly into gait analysis, we acquired motion capture data of a single healthy male (34 years, 95 kg) during flat ground walking and stair navigation. We received written informed consent from the volunteer under a study protocol approved by the University of Pennsylvania’s Institutional Review Board (Protocol # 824466). Reflective markers (9.5 mm, B&L Engineering, Santa Ana, CA, USA) were placed on the lower-extremities and tracked using motion capture. Markers were placed over a set of anatomic landmarks of the lower extremities to define a constrained musculoskeletal model: anterior and posterior superior iliac spines, medial and lateral knee condyles, medial and lateral ankle malleoli, calcaneus, and first and fifth metatarsal heads. An additional tracking marker was placed on both shanks and thighs, which we have shown to produce accurate kinematics [12]. The subject walked over ground at a self-selected speed (1.5 ± 0.1 m/s) and ascended and descended the staircase. We collected ten trials of each condition and processed the motion capture data using standard lab techniques [12]. To calculate internal joint moments during gait and stair navigation, we scaled a musculoskeletal model using subject-specific anthropometry and performed an inverse dynamics analysis [14]. We calculated the 95% confidence intervals [15] using both the uncorrected and corrected ground reaction forces during both over ground walking and stair navigation to visualize when uncorrected forces accounted for errors in joint kinetics.

## Results

The calibration procedure successfully reduced center of pressure errors in the force plates and both steps (**Table 1**, **Figure 3**). Before calibrating the force plates, the average center of pressure error (RMSE) was 6.7 mm and decreased to 3.3 mm after calibration. Despite being elevated 0.395 m from the force plates, the second step had similarly small errors compared to the force plates before (6.2 mm) and after (3.1 mm) the calibration procedure. The height of the fourth step (0.795 m) had larger effects on the center of pressure accuracy. Before calibrating the high step, the average center of pressure error was 19.6 mm but these errors decreased to 6.2 mm after the calibration procedure.

**Table 1.**
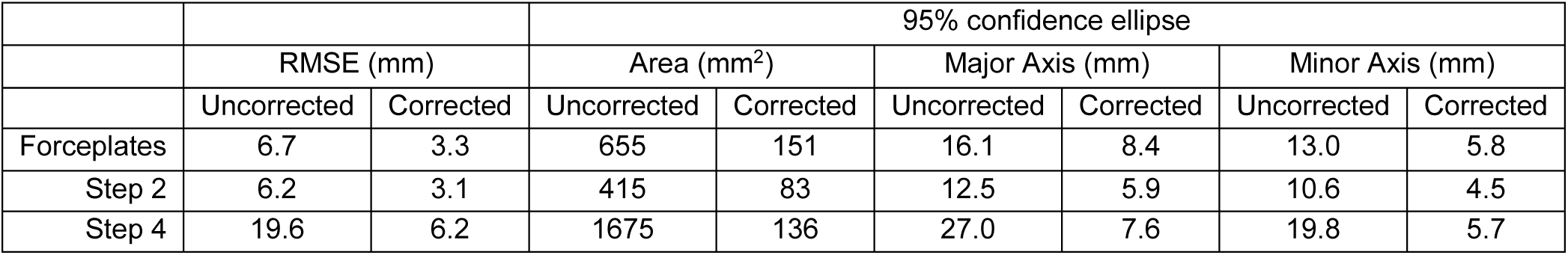
Uncorrected and corrected center of pressure errors

Lower extremity reaction moments were sensitive to the magnitude of ground reaction force errors and these errors were more pronounced at the proximal joints compared to the ankle (**Figures 4 and 5**). During level-ground walking, uncorrected ground reaction forces did not affect the calculated plantar flexion moments during stance. However, differences in the calculated knee and hip flexion moments during late stance were detected. Joint kinetics measured on the lower step (step 2) were not affected by using the uncorrected ground reaction forces in the inverse dynamics calculation. However, joint moments calculated from uncorrected ground reaction forces measured on the higher step (step 4) caused more consistent errors during both stair ascent and stair descent.

**Figure 4.**
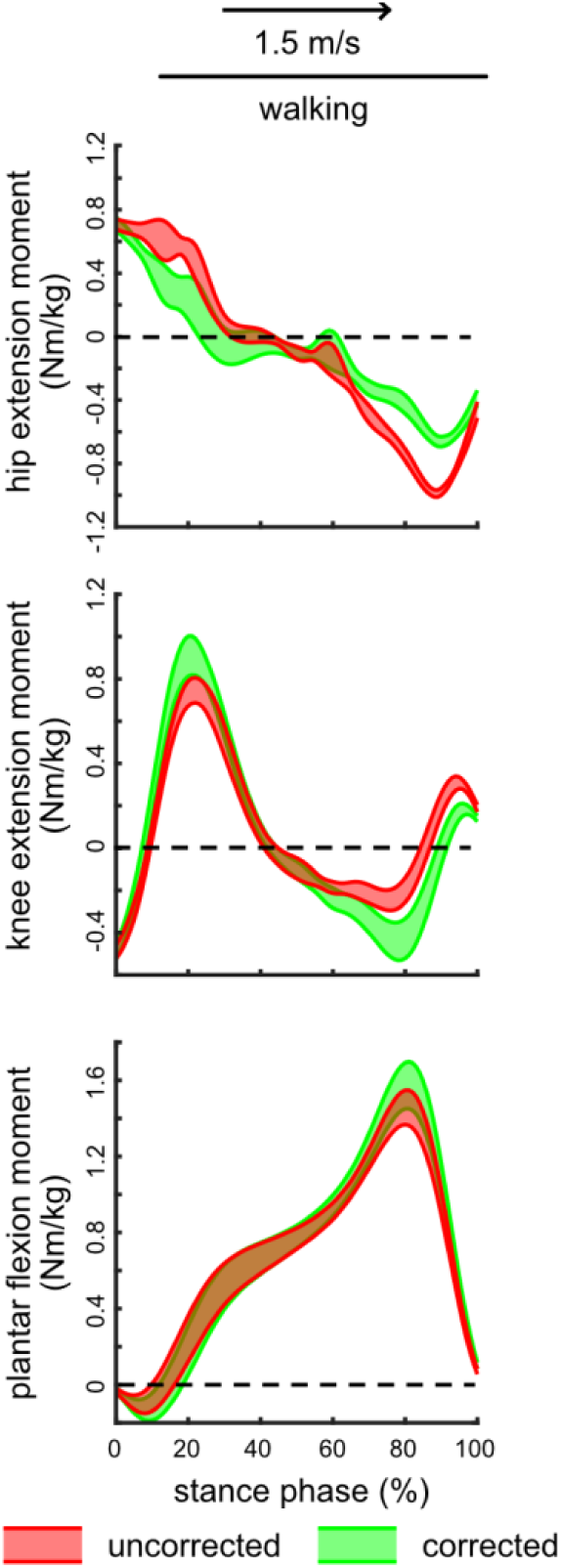
(1 column). Hip extension (*top*), knee extension (*middle*), and plantar flexion (*bottom*) internal moments during walking at 1.5 m/s were calculated using inverse dynamics with the uncorrected (*red*) and corrected (*green*) ground reaction forces. We analyzed 10 walking trials and calculated the 95% confidence intervals to visualize instances during stance where uncorrected ground reaction forces produce different joint moments compared to corrected ground reaction forces. Hip moments were more affected by uncorrected ground reaction forces than knee or plantar flexion moments. Dashed lines represent a transition from extension (+) to flexion (-) moments.

**Figure 5.**
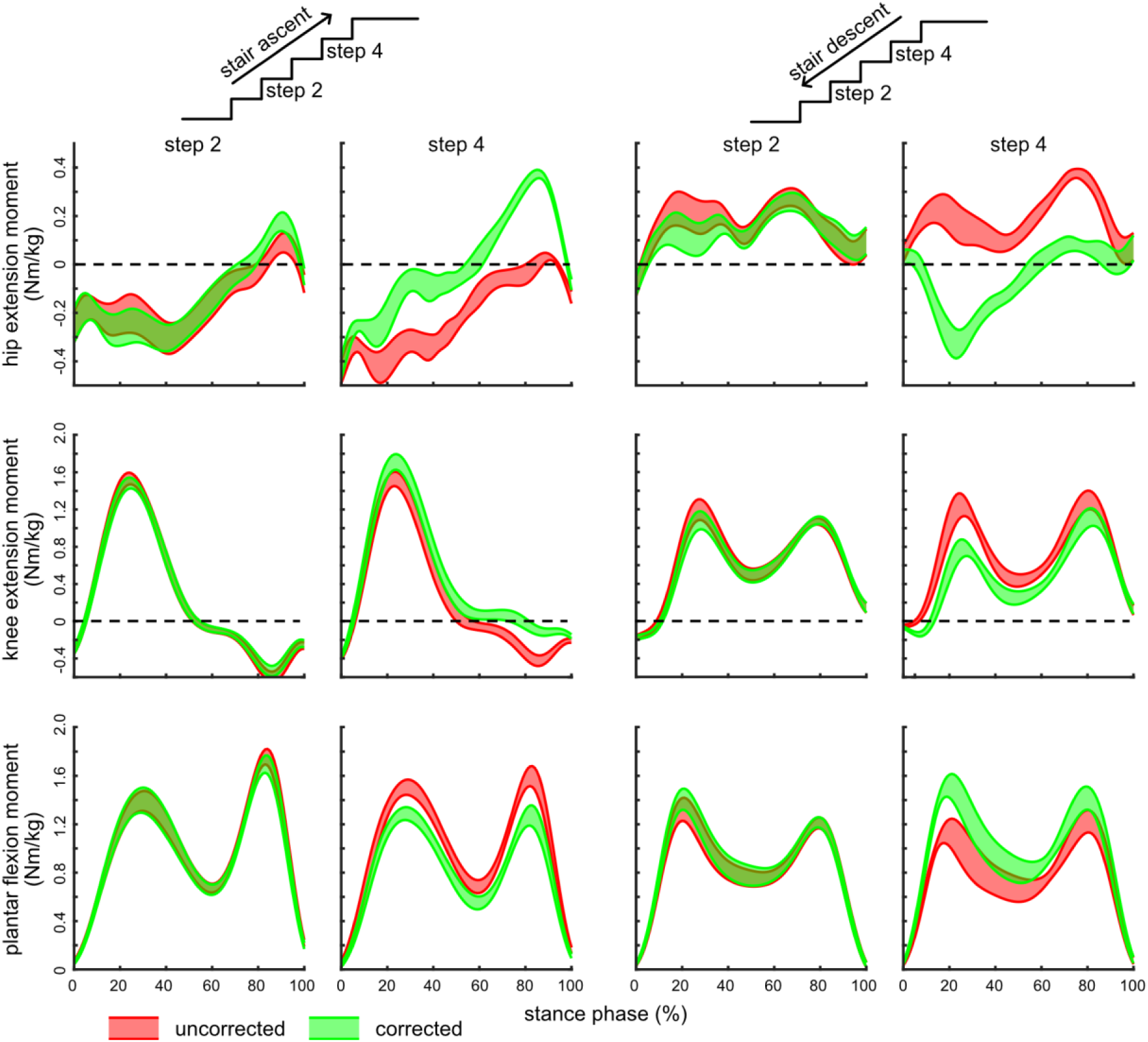
(2 column). Hip extension (*top*), knee extension (*middle*), and plantar flexion (*bottom*) internal moments during stair ascent (*columns 1 and 2*) and stair descent *columns 3 and 4*) were calculated using inverse dynamics with the uncorrected (*red*) and corrected (*green*) ground reaction forces. We analyzed 10 ascent and 10 descent trials and calculated the 95% confidence intervals to visualize instances during stance where uncorrected ground reaction forces produce different joint moments compared to corrected ground reaction forces. Lower extremity kinetics measured on the low step (step 2, *columns 1 and 3*) were not affected by uncorrected ground reaction forces. However, joint kinetics measured on the high step (step 4, *columns 2 and 4*) did differ – most notably at the hip – between the uncorrected and corrected ground reaction forces. Dashed lines represent a transition from extension (+) to flexion (-) moments.

These errors detected on the higher step (step 4) were consistent with the large anteriorly directed (when ascending) center of pressure errors that we were able to correct using the calibration procedure (**Figure 3**). These uncorrected center of pressure errors explained the over approximated plantar flexor moments and under approximated knee extension and hip extension moments during stair ascent. Conversely, these center of pressure errors resulted in similar errors with the opposite implications on joint kinetics with plantar flexor moments being under approximated and knee extension and hip extension moments being over approximated.

## Discussion

We designed and fabricated a staircase that leverages existing force plates embedded in our motion capture lab to assess joint kinetics during stair navigation. Based on our embedded force plates, we were able to position two steps on a set of three force plates positioned in a pyramid layout. Once the staircase assembly was positioned on the lab floor and force plates, we used a previously described calibration approach [11] to correct ground reaction forces applied to a rigid step assembly with two steps at known heights (**Figure 2**). These errors on the lower step (step 2) were small and similar in magnitude to uncorrected force plate measurements. However, the higher step (step 4) had much greater errors, mostly in the anterior direction, that were decreased by 68% following the calibration procedure. Our goal of this study was to develop a staircase that could easily be integrated into existing clinical gait analysis workflows and demonstrate high measurement fidelity. To this end, we included the complete bill of materials, mechanical drawings, experimental data, and calibration code we used in this project.

Our calibration protocol was based on a previous study that applied known loads to a force structure using an instrumented tool to calculate a calibration matrix using a least squares approach [11]. We first calibrated 3 embedded force plates and confirmed that center of pressure errors produced by our uncorrected force plates were similar in magnitude to those reported by Collins and colleagues. Despite resting on force plates, the loads applied to the low step (step 2) produced center of pressure errors that were similar in magnitude to the embedded force plates. These errors were reduced by 50% following the calibration procedure, which was similar in magnitude to the calibrated force plate measurements. However, increasing the step height to 0.795 m increased the uncorrected center of pressure errors by nearly 3-fold (**Figure 3**). While these errors were nearly half as large as the uncorrected center of pressure errors reported on an instrumented treadmill [11], these larger errors do show the limitations of measuring loads transferred through a wobbling mass. The errors in center of pressure had predictable effects on ankle, knee, and hip joint torques that were consistent with classic literature by McCay and DeVita [16]. Anteriorly shifted centers of pressure increased the calculated plantar flexor and hip extension moments while decreasing knee extension moments. (**Figure 5**). These small errors appear to have a greater impact on hip kinetics, in part to the smaller ground reaction force moment arms that translate to greater relative errors for the same absolute shift in center of pressure.

We developed an appreciation for several factors while designing and fabricating this staircase. Increasing the rigidity of the inner stair assembly is critical for both accurate measurements as well as recreating an authentic experience while navigating stairs. Our initial design used separate inner stair assemblies for the low (step 2) and high (step 4) steps. This layout resulted in a stable low step but a higher step that felt ‘unstable’ to the user. With this feedback during development, we secured these two inner steps to a common pair of aluminum bars that rested on the three embedded force plates. We also added 45 degree angled supports between the low and high steps to further increase stability.

This instrumented staircase paradigm could be easily modified to fit different force plate layouts. When considering step positioning on existing force plates, we found that the supporting foot pads of the step should be at least as long as the step is high. This decreases the likelihood that the ground reaction force passes outside of the base of support of the inner step. Because our force plates are 0.90 m wide, we did not experience any instability or step lift-off when applying medial-lateral loads. However, we did observe an asymmetrical error in the anterior-posterior errors of center of pressure on the higher step. This may have been because the applied loads were near the edge of the embedded force plate. To confirm the safety of these steps, we did two checks: 1) we applied anterior-posterior loads to the inner staircase assembly and confirmed that it did not move and 2) the inner staircase assembly has a small amount of separation with the outer assembly, which once in contact would serve as a mechanical stop for the inner assembly. We also added hand rails and 45 degree angled braces that increased the medial-lateral stability.

This mobile staircase can be effectively integrated into a clinical gait analysis workflow. Using an 18V cordless power drill, we are able to move and lower the staircase directly on top of the embedded force plates in less than 1 minute. Once in position, motion capture cameras mounted above eye level have largely unobstructed views of reflected markers placed on the lower extremities of test subjects. However, cameras on low tripods would likely have limited access to markers when positioned behind the elevated landing of the staircase. Once data is collected, we leveraged open-source musculoskeletal modeling software to perform the inverse kinematics and dynamic analyses (**Figures 4 and 5**) [14]. This approach provides a convenient visualizer to confirm force plate calibration and motion capture fidelity. Measuring ground reaction forces on two steps of different heights is necessary to quantify different joint loading patterns across a range of steps [17–19]. To our knowledge, this is the first staircase that rests on top of embedded force plates that measures ground reaction forces on high steps close to landings [20].

Several limitations affected this study. We designed this staircase specifically for our lab layout, which has a large continuous footprint of force plates (1.2 m by 0.9 m). Different configurations with smaller force plate footprints might limit the height and steps that can be measured with embedded force plates. Measuring joint kinetics on higher steps may be more sensitive to measurement errors (**Figure 5**). However, our calibration procedure reduced these errors nearly 3-fold, which may provide the necessary accuracy for studying specific components of stair navigation like transitioning to and from an elevated landing [17–19]. We decided to test this staircase paradigm on a single healthy adult to determine the isolated effects of using uncorrected and corrected ground reaction forces on gait and stair navigation joint kinetics. Our future work is aimed at understanding if patients with compromised plantar flexors who utilize compensatory mechanisms during other tasks [21,22] use similar strategies when navigating stairs.

In conclusion, our staircase design and calibration procedure provides researchers with a new tool to incorporate stair navigation biomechanics into their existing laboratory layout. Using an established force structure calibration procedure [11], we successfully reduced errors in the measured center of pressure applied using an instrumented tool to similar magnitudes previously reported in instrumented treadmill studies [11,13,23]. This staircase paradigm can easily be modified and adapted to meet the constraints of different laboratory layouts and workflows.

## Acknowledgments

We would like to thank Kaneel Senevirathne for assistance during pilot testing.

